# Neurocognitive patterns dissociating semantic processing from executive control are linked to more detailed off-task mental time travel

**DOI:** 10.1101/765073

**Authors:** Hao-Ting Wang, Nerissa Siu Ping Ho, Danilo Bzdok, Boris C. Bernhardt, Daniel S. Margulies, Elizabeth Jefferies, Jonathan Smallwood

**Affiliations:** Sackler Centre for Consciousness Science, University of Sussex, Brighton, United Kingdom; Department of Psychology, University of York, York, United Kingdom; Department of Psychiatry, Psychotherapy and Psychosomatics, RWTH Aachen University, Aachen, Germany; JARA, Translational Brain Medicine, Aachen, Germany; Parietal Team, INRIA, Neurospin, CEA Saclay, Gif-sur-Yvette, France; Mcconnell Brain Imaging Centre, Montreal Neurological Institute and Hospital, McGill University, Montreal, Canada; Frontlab, Institut du Cerveau et de la Moelle épinière, Paris, France

## Abstract

Features of ongoing experience are common across individuals and cultures. However, certain people express specific patterns to a greater extent than others. The current psychological theory assumes that individual differences in thought patterns occur because different types of experience depend on differences in associated neurocognitive mechanisms. Consequently, individual variation in the underlying neurocognitive architecture is hypothesised to determine the ease with which certain thought patterns are generated or maintained. Our study (N=178) tested this hypothesis using multivariate pattern analysis to infer shared variance among measures of cognitive function and neural organisation and examined whether these hidden structures explain reports of the patterns of on-going thoughts people experienced in the lab. We found that relatively better performance on tasks requiring primarily semantic knowledge, rather than executive control, was linked to a neural functional organisation that was associated, via meta-analysis, with task labels related to semantic associations (sentence processing, reading and verbal semantics). Variability of this functional mode predicted significant individual variation in the types of thoughts that individuals experienced in the laboratory: Neurocognitive patterns linked to better performance at tasks that required guidance from semantic representation, rather than those dependent on executive control, were associated with patterns of thought characterised by greater subjective detail and a focus on time periods other than the here and now. These relationships were consistent across different days and did not vary with task condition, indicating they are relatively stable features of an individual’s cognitive profile. Together these data confirm that individual variation in aspects of ongoing experience can be inferred from hidden neurocognitive architecture and demonstrate that performance trade-offs between executive control and long term semantic knowledge are linked to a person’s tendency to imagine situations that transcend the here and now.

## 2 INTRODUCTION

Ongoing experience is not always focused on the events, or tasks, taking place in the immediate environment. Sometimes we find ourselves absorbed by personally-relevant thoughts that are not directly related to events in the present, an experience that is often studied under the rubric of mind-wandering^1,2^. Psychological studies have established that there is substantial individual variation in self-generated experiences. Individuals whose experience is frequently directed away from the task in hand, perform poorly on tasks of sustained attention, reading and lecture comprehension, intelligence and working memory^3–5^. However, these individuals can perform better on tasks measuring creativity, problem-solving, or those that are semantic or episodic in nature^6–10^. Although we know that different types of ongoing experience make important contributions to our daily lives, we lack a clear understanding of the mechanisms that influence how and why this common everyday experience varies across people^11,12^.

Contemporary accounts assume individual differences in thought patterns occur in part because different types of experience depending on specific underlying neurocognitive mechanisms^13^. Consequently, individual variation in these underlying features will determine the ease with which certain thought patterns are generated or maintained. Consistent with this view, studies highlight attention and control systems as important in the prioritisation and maintenance of patterns of thought in a context-appropriate manner^14,15^. In contrast, activity patterns within regions allied to the default mode network (DMN), are associated with features of experience that are closely tied to the nature of the representations of the thoughts themselves^16–20^. In healthy participants, for example, changes in activity within the default mode network are linked to patterns of experience with rich subjective details in both episodic memory and working memory tasks contexts. In patients with forms of dementia that target elements of the default mode network (including posterior cingulate cortex), patterns of off-task experience are less intense^21^ and scene construction is impaired, particularly global features of the experience^22^. Likewise, patients with semantic dementia, in which atrophy is focussed on the anterior temporal lobes, have deficits in the ability to imagine the future^23^. Finally, lesions to the hippocampus impact on the episodic content that often occupies the absent mind^24^.

The complex associations between aspects of ongoing experience, measures of cognition and neural function suggests that individual variation in specific patterns of experience may be reflected by composite features, encompassing differences in both cognition and neural function. For example, according to contemporary views of mind-wandering^11,13^ a chronic tendency towards off-task thoughts may depend on a relative weakness in the ability to perform well on tasks of executive control, relative strengths on the ability to generate cognition based on memory representations^13,25^, or a combination of both. Moreover, these changes could be mirrored by equally complex neural patterns (e.g. See prior study^26^). At first glance, this heterogeneity of functional association may seem to undermine attempts to understand the individual differences that underpin different aspects of ongoing thought. However, in our study, we take advantage of this richness by using Sparse Canonical Correlation Analysis (SCCA)^27,28^, which capitalises on machine learning and multivariate data patterns to infer the underlying dimensional structure that best explains complex data matrices.

Our prior studies^26,29^ used SCCA to infer hidden patterns in resting-state data that are linked to individual variation in the patterns of thoughts themselves. In the current study, we focus on whether composite neurocognitive features that describe variation in different types of cognitive process (e.g. executive control, semantic memory) and neural organisation describe meaningful trait variation in patterns of ongoing thoughts. We recorded performance in a large cohort of participants on a battery of tasks, including those more reliant on semantic processing (e.g. Graded naming, semantic association) and those with a higher reliance on executive control (e.g. Digit span and task switching). In the same participants, we acquired measures of both structural and functional brain organisation. In addition, the same participants took part in a multi-day study in which they performed tasks that varied in their working memory demands, to test for context-dependent effects on experience. While participants performed these tasks, we used multi-dimensional experience sampling (MDES), which requires participants to provide self-reported descriptions of their experience. This method asks participants on many occasions to provide reports on multiple questions that describe the content and form of their ongoing thoughts.

In our investigation, we applied multivariate CCA to the sets of behavioural and brain data to generate a set of modes that reflect joint descriptions of the data from both domains (i.e. Dimensions that reflect commonalities between how a person performed on the tasks and their brain organization at rest). We used this set of neurocognitive descriptions to predict patterns of ongoing experience recorded in the laboratory to understand if these neurocognitive modes were related to the experiences that participants reported in the laboratory. We aimed to test the hypothesis that individual variation in the tendency to be absorbed in thoughts related to other times and places, reflects a trade-off between neurocognitive processes that rely on more generative aspects of cognition and those more closely associated with greater executive control^13^.

## 3 RESULTS

### 3.1 IDENTIFICATION OF NEUROCOGNITIVE MODES OF VARIATION

Our current study aims to understand whether the composite neurocognitive features are linked to the individual differences in ongoing thoughts. We first identified neurocognitive hidden structures between cognitive performance and whole-brain functional connectivity as the two input matrices. The application of CCA to the neural and behavioural data identified two reliable modes that reached statistical significance based on permutation testing (See Methods). A meta-analysis of the functional connectivity summary was performed using Neurosynth^30^ to facilitate our interpretation of these whole-brain patterns (see Methods). These cognitive and neural patterns are presented in Figure 1. The cognitive patterns and the meta-analysis results are presented as world clouds. The size of the word indicates the strength of the relationship and the colour of the direction of the association.

**Figure 1.**
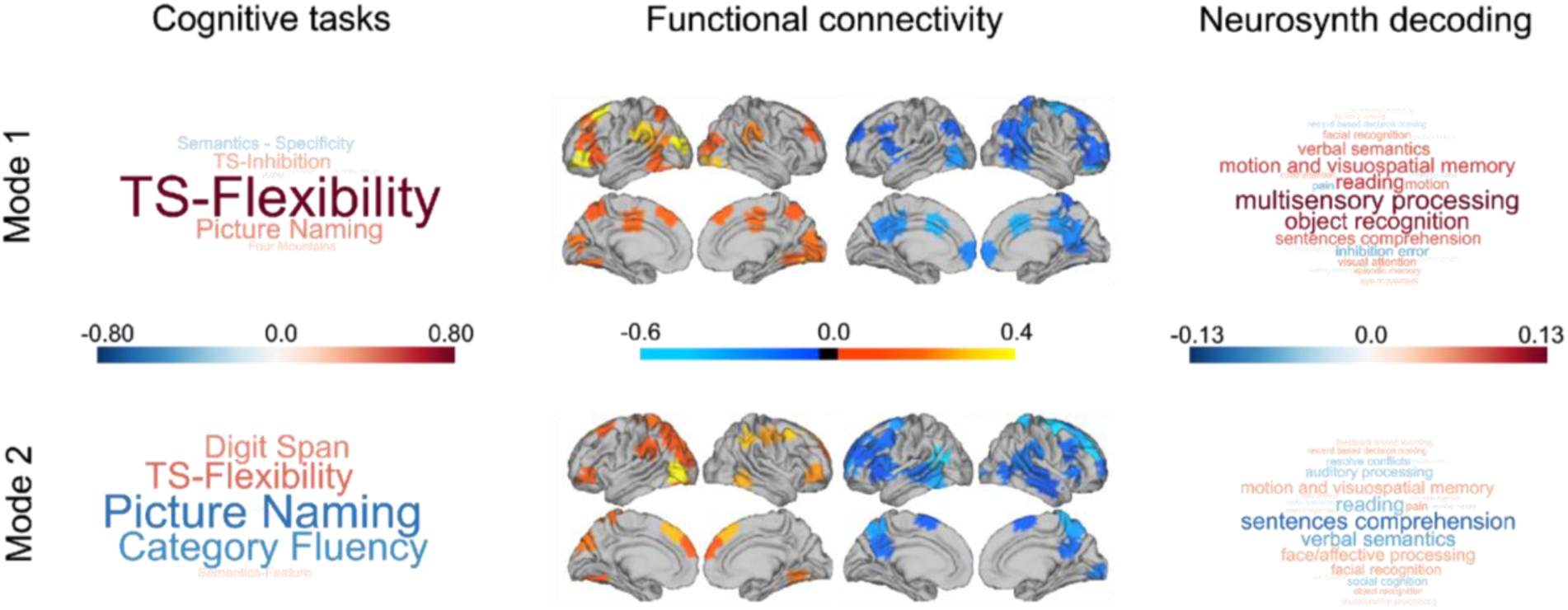
Two reliable neurocognitive modes uncovered through CCA. In both sets of word clouds, the size of the word indicates the strength of the relationship and the colour the direction of the association. The left-hand panel shows the task loadings for each mode in the form of a word cloud. The central panel presents the brain regions that contributed the most to each mode. The orange-yellow maps present the positively connected nodes and the blue maps show the negatively connected modes. The right-hand panel shows word clouds that describe the results of a quantitative meta-analysis of this functional data performed using Neurosynth (see Methods).

Behaviourally, mode 1 (r = 0.57, p = 0.036) is reflected by positive loadings on task switching (both flexibility and inhibition) and picture naming, with relatively weak negative loadings on the specific feature matching and strong semantic association task. These two tasks both required the ability to access semantic knowledge but not flexibility in semantic processing^31^. In neural terms, functional connectivity was dominated at the positive end of the dimension by lateral visual, the left temporoparietal junction and the left ventral lateral prefrontal cortex. In contrast, the negative end of the functional dimension was dominated by a region of middle cingulate, the anterior motor cortex and lateral occipital cortex. Meta-analysis highlighted positive loadings for terms linked to perceptual processing, including “object recognition”, “motion and visuospatial memory” as well as terms linked to semantic processing (“verbal semantics” and “sentence comprehension”). It is noteworthy that the task loading most heavily on the behavioural features of this dimension involved switching between judgements based on shape, movement and colour. This pattern is broadly reflected in the meta-analysis of the neural data (“motion and visual-spatial processing”) suggesting reasonable convergence between the brain and behavioural descriptions of the mode.

Mode 2 (r = 0.59, p = 0.011) was associated with strong positive loading on tasks that require executive control (“digit span” and “flexibility”) and strong negative loadings on tasks that required self-generation of overt responses based on semantic knowledge (“picture naming” and “category fluency”). The neural patterns emphasised regions of the ventrolateral visual cortex as well as the pre-supplementary area in yellow, while the negative end in blue was dominated by the left angular gyrus, the dorsal precuneus and the dorsomedial prefrontal cortex. A meta-analysis of the functional data highlighted a clustering of terms associated with semantic tasks at the end associated with better picture naming and category fluency (“sentence comprehension”, “verbal semantics” and “reading”). Similar to mode 1, therefore, there was a mapping between the pattern of behaviour for mode 2 (better performance on semantic tasks) and meta-analysis of the associated neural patterns (a relative preference for terms with semantic links).

### 3.2 TRADE-OFFS BETWEEN SEMANTIC AND EXECUTIVE CONTROL PREDICT DETAILED OFF-TASK MENTAL TIME TRAVEL

Having determined two reliable neurocognitive modes, we next tested the hypothesis that these explained significant variance in the data collected in the experience sampling phase of the experiment. We conducted a multiple multivariate regression in which the average responses to the experience sampling questions were the dependent variables. We included scores for each individual for each mode of brain-behaviour co-variation as explanatory variables. Age, gender and mean frame displacement were included as nuisance covariates. This analysis revealed that the variation in mode 2 was associated with the experience sampling questions at the multivariate level (F(164, 13)=2.87, p=0.009, *η*^2^=0.19). No similar association was seen with mode 1 (F(164, 13)=0.64, p=0.82, *η*^2^=0.05). Figure 2B illustrates this multivariate association between mode 2 and reports of ongoing experience in the form of a word cloud in which the size and colour of the items reflect the strength and direction of the relationship. It can be seen that individuals who tended to be better at tasks that depended on semantic knowledge to generate task responses (see Figure 2C) tended to emphasise a pattern of thoughts high on subjective detail, and that were unrelated to the task, and focused on time periods other than the present.

**Figure 2.**
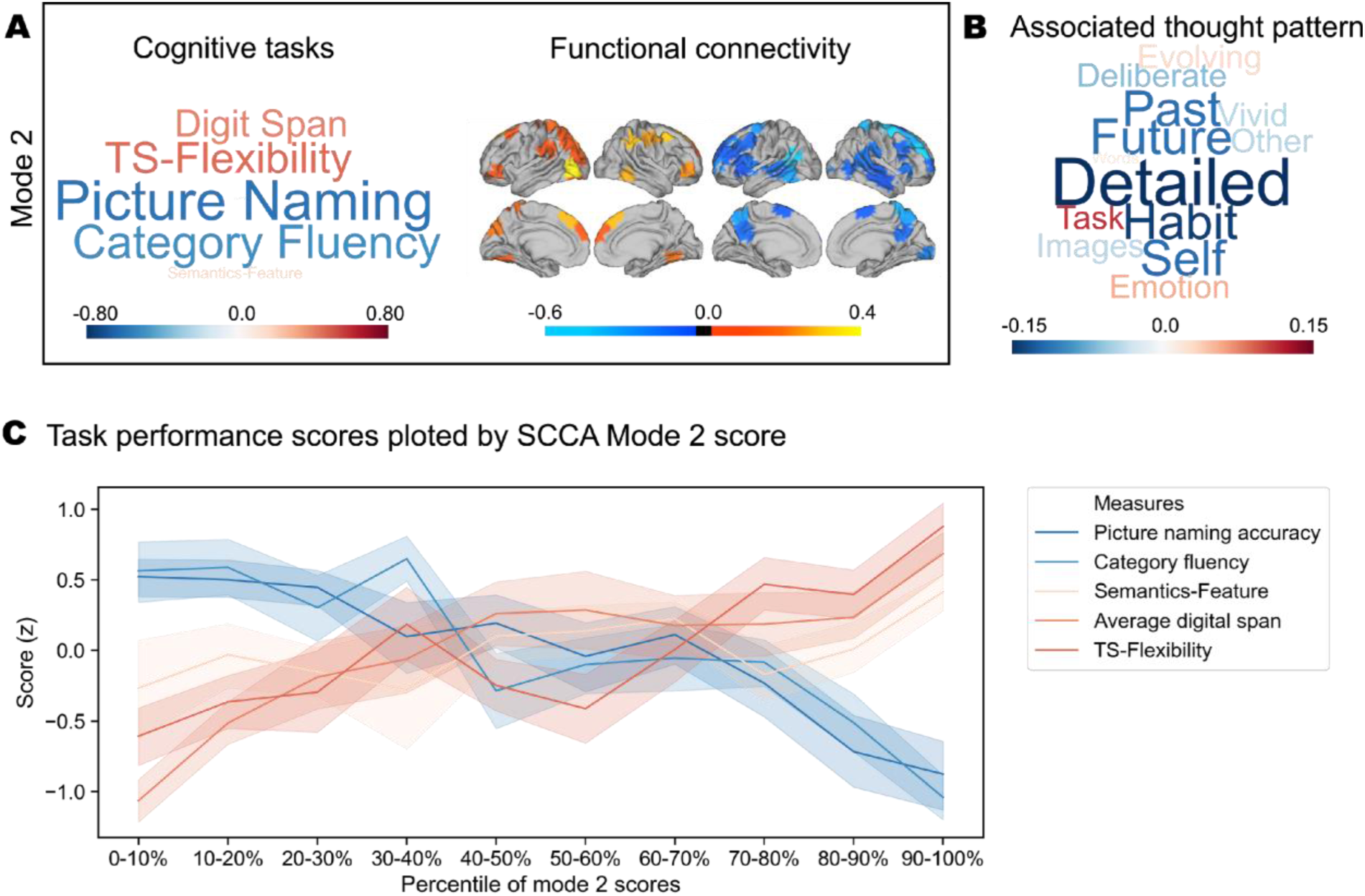
The patterns in the experience sampling questions associated with mode 2. (A) The neurocognitive associations that constitute mode 2; (B) The multivariate relationship between mode 2 and patterns of ongoing thought displayed in the form of a word cloud in which the size and colour of the items reflect the strength and direction of the relationship. Individuals who tended to be better at tasks dependent on semantic knowledge to generate task responses tended to endorse patterns of thoughts high on subjective detail, and that were unrelated to the task, and focused on other periods (past and future). (C) Standardised task performance scores plotted alone the percentile of mode 2 score. The ribbon plot shows that mode 2 captures patterns of individual differences in the trade-offs between executive function and semantic knowledge. The shaded bars describe the 95% confidence intervals of the mean.

### 3.3 THE RELATIONSHIP BETWEEN NEUROCOGNITIVE MODES ARE STABLE ACROSS TASKS CONDITIONS AND DAYS

Having determined an association between neurocognitive modes of variation and patterns of ongoing thought, we next examined how stable this relationship was across task context and day. In our experience sampling study, participants performed alternating task blocks of lower (0-back) and higher (1-back) working memory demands. If the relationship between the functional modes and patterns of ongoing thoughts are context-specific, then they could change across the conditions or days of the experiment. We examined whether similar patterns of thoughts emerged across these contexts by running separate multiple multivariate linear regressions on each task condition. The outcome of these analyses is presented in Figure 3. In Figure 3A and 3B, it can be seen that this produced similar patterns of experience in both tasks, indicating that the association with mode 2 was robust across task contexts (1-back: F(164, 13)=2.97, p=0.0006, *η*^2^=0.19; 0-back: F(164, 13)=2.43, p=0.0048, *η*^2^=0.16; range of correlation across task contexts [0.93 0.98] with all p<0.001). Importantly, the patterns produced by these analyses were highly correlated with each other, and with the pattern produced by the overall analyses. Next, we examined whether these relationships are consistent across different days of our experience sampling study. We carried out three multiple multivariate regressions in which average scores for each day were dependent variables. The results of this analysis is presented in Figure 3C and 3D where it can be seen that the relationship remained relatively stable across the sessions of our experiment (Day 1: F(164, 13)=1.91, p=0.0327, *η*^2^=0.13; Day 2: F(161, 13)=1.62, p=0.0855, *η*^2^=0.12; Day 3: F(129, 13)=2.93, p=0.0009, *η*^2^=0.23; range of correlation among days [0.83 0.96] with all p<0.001). As with the analysis of different task contexts, we found that the patterns of thoughts produced by the analysis on each day were consistent with each other and also with the pattern produced by the overall analyses. It is important to note that the sample size for each day varies because some participants did not participate in all sessions.

**Figure 3.**
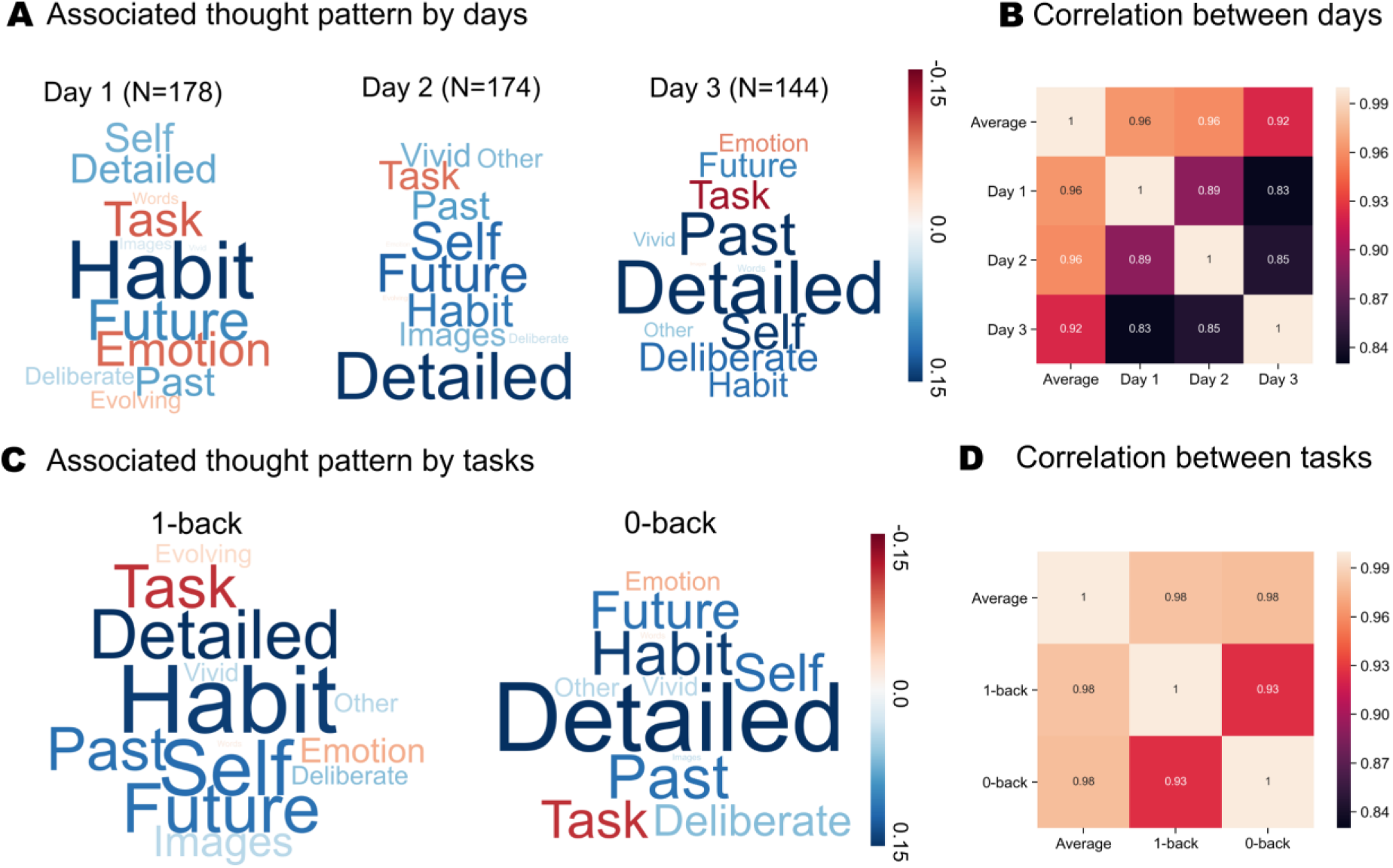
The stability of mode 2 in predicting thought patterns across task context and day. (A) Similar multivariate associations between mode 2 and the patterns of ongoing thought emerged in both by tasks conditions; (B) correlations between the patterns of ongoing thought observed in general and across both tasks; all p-value < 0.001; (C) Similar patterns of ongoing thought were observed across each day of the study; (D) correlations between the patterns of ongoing thought observed in general and across each day of the study; all p-value < 0.001.

### 3.4 TRADE-OFFS BETWEEN SEMANTIC AND EXECUTIVE CONTROL AND THEIR RELATION TO THE CORTICAL STRUCTURE

Our last set of analyses considers the relationship between the two modes and a more trait-like element of cortical organisation: whole-brain grey matter thickness. Cortical thickness is a relatively stable feature of an individual’s neuroanatomy and is related to genetic factors^32,33^, personality traits, and neuropsychiatric disorders (e.g. ^34–36^). If we found associations between the individual variation in a mode and individual differences in cortical grey matter structure, this would enhance our confidence that our CCA mode described a reasonably stable neurocognitive trait.

To understand the association between variation in the two functional modes, and variation in the thickness of cortical grey matter, we conducted a regression in which a matrix describing thickness at each vertex for each individual were the dependent variables. The average score for each individual on each mode was included as explanatory variables. Age and gender were both entered as between participant explanatory variables. This initial model yielded no significant results.

Next, we conducted a more exploratory analysis in which each individual’s score for the behavioural and neural aspects was entered in the same model as separate explanatory variables (a total of 4 evs). This analysis revealed significant associations (FWE-corrected p < .05, uncorrected-p < .0025) with both the behavioural and neural aspects of mode 2 and are presented in Figure 4. The behavioural dimension, describing performance trade-offs between tasks tapping semantic and executive domains, was with variation in cortical thickness in a region of subgenual anterior cingulate (mode 2 CT: r = -.29, p < .001). Thickness in this region was higher for individuals better at semantic than executive tasks. Also, the functional dimension was associated with cortical thickness in a region of motor cortex immediately posterior to the central sulcus (mode 2 FC: r = .26, p < .001). Thickness in this region was smaller towards the semantic end of this dimension.

**Figure 4.**
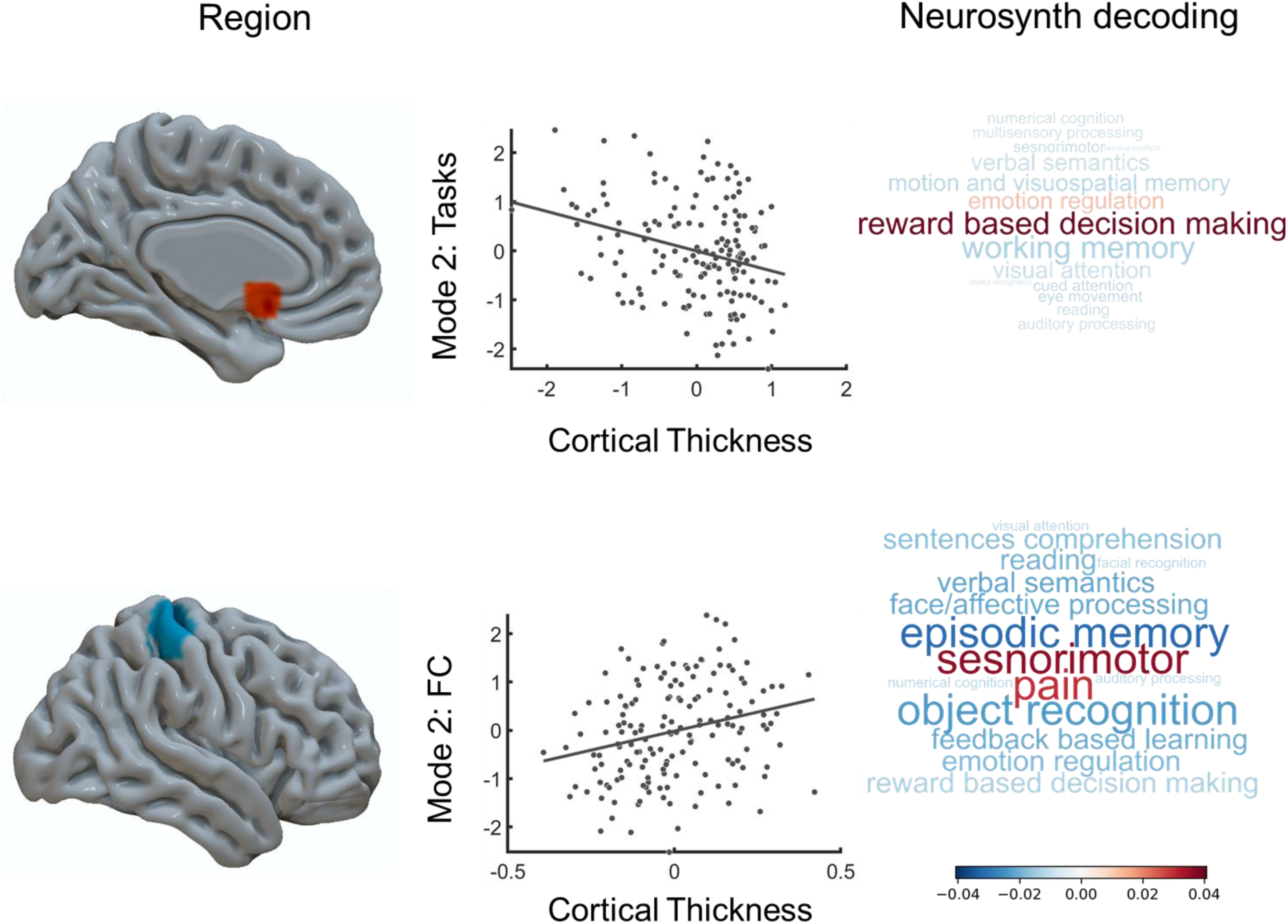
Exploratory cortical thickness analysis. We used a general linear model to explore whether individual variation in the canonical modes was also reflected in the structural organisation of the cortex. We found that the task component in mode 2 is related to the variation in a region of subgenual anterior cingulate among. Thickness in this region was higher for individuals better at semantic than executive tasks. The functional connectivity component of mode 2 was associated with cortical thickness in a region of motor cortex immediately posterior to the central sulcus. Thickness in this region was smaller towards the semantic end of this dimension.

### 3.5 SUMMARY

Using machine learning, we found that relatively better performance on tasks requiring the generation of responses based on semantic knowledge than executive control was linked to a pattern of functional organisation that associated via meta-analysis processes with semantic features (sentence processing, reading and verbal semantics). Features of this functional mode predicted significant variation in the types of thoughts that individuals experienced in the laboratory: Performing better at tasks in which responses were guided by semantic representations, than those dependent on executive control, was associated with patterns of thought characterised by greater subjective detail and a focus on time periods other than the here and now. Finally, a cortical thickness analysis identified structural correlates of this functional mode, suggesting that this mode is a reasonably stable neurocognitive trait.

## 4 DISCUSSION

Our study aimed to understand why certain people spontaneously engaging more in certain patterns of thoughts than do others. We focused on testing the prediction of contemporary accounts of ongoing thought, which assume that different aspects of ongoing experience are related to different underlying neurocognitive processes^13,37^. Accordingly, the ease with which an individual engages in particular forms of ongoing thought should be related to the degree to which their neurocognitive architecture emphasises the relevant underlying neurocognitive process. In particular, we were interested in testing the broad view that periods of imaginative thinking during states of mind-wandering rely more on the ability to self-generate information from long term memory than do thoughts focused on the present^13^.

Consistent with this view, we found that individuals who emphasise patterns of thoughts about other times and places with rich subjective details, rather than the task in hand, show evidence of a trade-off towards expertise at semantic processing at the cost of executive control. Behaviourally, ‘detailed mental time travel’ was linked to better performance on tasks that rely on generating a response based on semantic knowledge rather than those that tap executive control. A meta-analysis of functional connectivity of the neural component of mode 2 suggested an association with patterns seen in tasks that rely on semantic processing (such as verbal semantics and sentence processing). These data provide converging evidence from both brain and behaviour, that the tendency to generate thoughts that extend beyond the current moment is linked to a more general ability to generate cognition based on existing semantic knowledge. Our finding confirms predictions theoretical accounts that assume that patterns of off-task thought are partly generated using information from long-term memory^13^. This has several implications for our understanding of the underlying mechanisms that give rise to different types of experience.

First, our study adds to a growing body of work showing that semantic processes play a key role in aspects of human imagination. Studies of patients with semantic dementia have shown that they have deficits in future thinking, as well as the inability to represent detailed scenes in imagination^23^. Moreover, dementia that results in atrophy to the posterior cingulate cortex impacts upon the intensity of mind-wandering episodes^21^ and our study highlights the connectivity of this region as important for patterns of mental time travel with vivid detail. Our study, therefore, builds on prior work that uses lesion based methods by showing that in a young healthy population individuals who perform better on task reliant on semantic processes, tend to report more detailed mental time travel. Together these studies provide converging support for a neural architecture that is implicated in tasks involving semantic processing as playing a role in imagination and in particular, content, level of detail, or perhaps both.

Second, our study has important methodological implications for contemporary accounts of ongoing thoughts. For example, work on mind-wandering has found that this pattern of experience has both costs and benefits in daily life. Deleterious aspects associated with excessive mind-wandering have been linked to problems in executive control and links to affective difficulties, while advantages have been linked to better performance on tasks that rely on the generation of information (creativity, semantic processing and self-reference)^38^. Our study suggests that patterns of reports that correspond broadly to contemporary views of the mind-wandering state (i.e. ^13^) can be understood as the consequence of trade-offs between different underlying qualities of cognition. In this way our highlights that certain questions regarding specific mental states can be understood as emergent properties from interactions within a more general component process architecture. One advantage of our approach is that it produces categories of experience in a data-driven manner, and so circumvents definitional questions regarding what constitutes a specific mental state^2,39,40^. There are, however, some important limitations that should be borne in mind when considering our study. We used a battery of tasks that primarily operationalised self-generated processing through the lens of tasks tapping semantic processing and creativity. This decision was pragmatic: such tasks provide a reasonably direct indication of one aspect of self-generated processing because they require individuals to use conceptual knowledge to guide behaviour. It was also well aligned with the expertise of our lab. This design choice, however, places an important boundary condition on the interpretations that should be placed on our findings. Our study highlights a robust association between how we use knowledge of the world around us to generate behaviour and the patterns of thoughts we generate in imagination. As we did not collect other aspects of self-generated processing from the participants (such as autobiographical memory or emotional processing), it remains an open question what role individual differences in these processes play in patterns of ongoing thought. It will be important for future work to extend our methodological approach to deepen understanding how variation in the emotional, or episodic domain contribute to patterns of ongoing thoughts, and how similar or different these are to variation in semantic knowledge. It is also important to note that our study focused on patterns of ongoing through the lens of a trait. There are important aspects of patterns of ongoing thought that can only be properly understood when it is recorded simultaneously with ongoing neural activity^41^. It is important to consider how these individual differences translate into momentary patterns of neural activity. In this context, it is worth noting that we have recently shown that that neural signals in both the dorsal left pre-frontal cortex, and the posterior cortex are linked to momentary differences in off-task content, and levels of subject detail respectively^15^. These regions are both highlighted by our analysis as members of the neural network linked to semantic processing, and to reports of experiences which emphasise off-task episodes with a high degree of subject detail (See also Kam and colleagues^42^ for a conceptually similar conclusion). This association provides reasonable grounds to suggest that it could be profitable for future work to explore how patterns of neurocognitive traits at rest relate to the momentary neural patterns observed when the same pattern of thought occurs.

In conclusion, our study suggests that the reason why certain people engage more in certain patterns of thoughts than do others is related to the emphasis their neurocognitive architecture places on different types of process. We found experiences that emphasise detailed mental time travel, are linked to neural indices indicative of relative expertise in tasks reliant on semantic processing. Our study, therefore, underlines the core role that the way we understand the world around us plays in what we construct using imagination. It is worth noting that that occurrence of imaginative thoughts are common across cultures^43^ and have links to both beneficial and detrimental aspects of wellbeing^38^. Accordingly, it seems possible that the structure and ease with which conceptual knowledge can be mobilised in imagination will be an important influence on how a ubiquitous aspect of daily life impacts upon a person’s happiness and success.

## 5 METHOD

### 5.1 PARTICIPANTS

Two hundred and seven healthy participants were recruited from the University of York (132 females, 75 males; age range = 18-31 years, M = 20.21, SD = 2.36). This analysis included two data sets with some shared measurements and the same MRI protocol. Participants were right-handed native English speakers with normal or corrected-to-normal vision and no history of psychiatric or neurological illness. Participants underwent MRI scanning, completed a 1-hr online questionnaire. Of the first cohort, participants attended three (165 participants; 99 females, 66 males; age range = 18-31 years, M = 20.43, SD = 2.63) 2-hr behavioural testing sessions to complete a battery of cognitive tasks. The second cohort (42 participants; 33 females, 9 males; age range = 18-23 years, M = 19.79, SD = 1.37) underwent two 2-hr behavioural testing sessions to complete a battery of cognitive tasks. The behavioural sessions took place within a week of the scan. Twenty-nine participants were excluded from the multivariate pattern analysis because they failed to complete all of the behavioural testing sessions or failed the imaging data quality check. In total, 178 participants (113 females, 65 males; age range = 18-25 years, M = 19.81, SD = 1.66) were included in the multivariate pattern analysis and the comparison with cognitive performance. Participants were rewarded with either a payment of £10 per hour or a commensurate amount of course credit. All participants provided written consent prior to the fMRI session and the first behavioural testing session. Approval for the study was obtained from the ethics committee of the University of York Department of Psychology and the University of York Neuroimaging Centre.

### 5.2 MRI ACQUISITION

Structural and functional data were acquired using a 3T hdx excite MRI scanner (GE Healthcare, Little Chalfont, UK) utilising an eight-channel phased-array head coil tuned to 127.4 MHz at the York Neuroimaging Centre, University of York. Structural MRI acquisition in all participants was based on a T1-weighted 3-D fast-spoiled gradient-echo sequence repetition time (TR) = 7.8 s, echo time (TE) = minimum full, flip angle = 20°, matrix size = 256 x 256, 176 slices, voxel size = 1.13 x 1.13 x 1 mm^3^. Resting-state activity was recorded from the whole-brain using single-shot 2-d gradient-echo-planar imaging TR = 3 s, TE = minimum full, flip angle = 90°, matrix size = 64 x 64, 60 slices, voxel size = 3 x 3 x 3mm^3^, 180 volumes. Participants viewed a fixation cross for the duration of the 9-min fMRI resting-state scan. A fluid-attenuated inversion-recovery (FLAIR) scan with the same orientation as the functional scans were collected to improve co-registration between subject-specific structural and functional scans.

### 5.3 RESTING-STATE DATA PREPROCESSING

All preprocessing and denoising steps for the functional MRI data were carried out using the SPM software package (Version 12.0) and Conn functional connectivity toolbox (Version 17.f), based on the MATLAB platform (Version 17.a). The first three functional volumes were removed to achieve steady-state magnetisation. The remaining data were first corrected for motion using six degrees of freedom (x, y, z translations and rotations), and adjusted for differences in slice-time. Subsequently, the high-resolution structural images were co-registered to the mean functional image via rigid-body transformation, segmented into grey matter, white matter and cerebrospinal fluid probability maps, and all functional volumes were spatially normalized to Montreal Neurological Institute (MNI) space using the segmented images and a priori templates. This indirect procedure utilizes the unified segmentation normalization framework, which combines tissue segmentation, bias correction, and spatial normalization in a single unified model. No smoothing was employed, complying with recent studies that report the negative influence of this procedure on the construction of connectivity matrices analysis.

Moreover, a growing body of literature indicates the potential influence of participant motion inside the scanner on the subsequent estimates of functional connectivity. To ensure that motion and other artefacts did not confound our data, we have employed extensive motion-correction and denoising procedures, comparable to those reported in the literature. In addition to the removal of six realignment parameters and their second-order derivatives using the general linear model (GLM), a linear detrending term was applied as well as the CompCor method that removed five principal components of the signal from white matter and cerebrospinal fluid. Moreover, the volumes affected by motion were identified and scrubbed based on the conservative settings of motion greater than 0.5 mm and global signal changes larger than z = 3. Though recent reports suggest the ability of global signal regression to account for head motion, it is also known to introduce spurious anti-correlations and was thus not utilised in our analysis. Finally, a band-pass filter between 0.009 Hz and 0.08 Hz was employed to focus on low-frequency fluctuations.

The Craddock connectivity-based parcellation^44^ was selected as the full brain parcellation. We used a set of 100 regions (K = 10) from the two-level spatial connectivity-based parcellation. The Craddock atlas clusters delineate spatially coherent regions that are more functionally homogenous, thus it demonstrated good dimension reduction ability in the context of resting-state functional connectivity analysis. The spatial connectivity-based parcellation focuses on similarity between functional connectivity maps^44^. Fully connected, undirected and weighted matrices of bivariate correlation coefficients (Pearson’s r) were constructed for each participant using the average BOLD signal time series obtained from all the 100 ROIs described above. The off-diagonal of each correlation matrix contained 4950 unique measures of region-region connection strengths (i.e., the upper or lower triangle of the network covariance matrix). This approach provided a measure of connection strength of the whole-brain for each participant. Finally, Fisher’s r-to-z transformation was applied to each network covariance matrix.

### 5.4 BEHAVIOURAL DATA

#### 5.4.1 Cognitive tasks

We selected 9 cognitive tasks that are common across the two cohorts. The selected tasks measure cognitive functions that have been examined in previous mind-wandering literature, encompassing executive control (digit span, task switching task^45^), generation of information (unusual uses task^46^, verbal fluency task), semantic memory (semantics picture-word matching tasks^47^: relatedness judgement tasks and feature matching task^48^), episodic memory (paired-associate task^49^, four mountains task^50^), and fluid intelligence (Raven Advanced Progressive Matrices; RAPM^51^).

Thirteen cognitive scores were calculated from the selected tasks. Performance of the digit span task was represented as the average of digit span in the forward and backward recall conditions. The verbal fluency score is the contrast of the category condition and letter condition (category - letter). The previous research^31^ shows that category fluency is more dependent on semantic memory, while letter fluency is more executively demanding. Picture naming tasks, the four mountains tasks, RAMP were summarised with accuracy scores. The task switching measure provided two scores (a) flexibility^*^ as the ability to switch from a different condition and (b) inhibition as the ability to suppress information from the previous trial. The calculation of the task switching contrast can be found in the original study^43^. All the semantics related judgement tasks, feature matching task, and the paired-associate task were summarised using efficiency scores. The efficiency scores were calculated as reaction time divided by accuracy. A smaller score indicates better performance, thus the scores were reversed to ease the interpretation. In the semantics picture-word matching task^47^, a picture probe was matched either to an associated word (for strong, weak and word conditions) or an associated picture (in the picture condition) or to the name of the item at a superordinate or specific level. We calculated three contrasts based on the semantics modules tested: (a) strength (strong – weak), modality (picture – word), and (c) specificity (specific – general). All the scores were brought to a common scale by variance scaling to 1 and mean centring to 0 for the subsequent analysis.

#### 5.4.2 Experience sampling

We assessed the contents of experience in the context of a simple task that manipulated working memory load using a block design (see prior published examples of this task^7,52^). This task was performed at the beginning of each laboratory session to minimize the contribution of participant fatigue to these experiential measures. Measuring experience over multiple days provided us with a more comprehensive description of participants’ ongoing thought at the level of a trait than would have been possible in a single experimental session.

In both conditions, non-target trials involved the presentation of pairs of shapes appearing on the screen divided by a vertical line. The pairs could be a circle and a square, a circle and a triangle, or a square and a triangle for six possible pairs (two different left/right configurations for each). The pairs never had shapes of the same kind (e.g. A square and a square). In both tasks, following an unpredictable sequence of non-target trials, a target trial was presented in which participants had to make a manual response. The target was a small stimulus presented in either blue or red across conditions, with the colour counterbalanced across participants. In the 0-back condition, two shapes flanked the target and the target would be identical to one of the two flanking shapes. Participants had to indicate by pressing the appropriate button which shape matched the target shape. In the 1-back condition, the target was flanked by two question marks and participants had to respond depending on which side the target shape was on the prior trial. Responses were made using the left and right arrow keys. Fixation crosses presentation ranged from 1.3–1.7 seconds in steps of 0.05 seconds, non-targets were varied from 0.8–1.2 seconds in steps of 0.05 seconds. Targets always ranged from 2.1–2.5 seconds in steps of 0.05 seconds and a response from participants did not end the target presentation.

Multi-dimensional experience sampling (MDES) was used to describe experience during the 0-back/1-back task. This technique uses self-report to assess the contents of experiences on a number of dimensions. The thought probes first asked participants to rate their level of task focus (“My thoughts were focused on the task I was performing.”) on a sliding scale from 0 (completely off-task) to 1 (completely on task). Participants then answered 12 randomly presented questions regarding the content and form of their experience just before they were probed. These questions (described in Table 1) were based on prior studies adopting this approach to measure self-generated thought^7^. There was a 20% chance of a thought probe being presented instead of a target with a maximum of one thought probe per condition block of 0-back and 1-back. In each session, an average of 14.07 (SD = 3.30, range 6 – 25) MDES probes occurred; in the 0-back condition an average of 7.02 (SD = 2.36, range 2 – 14) MDES probes occurred and in the 1-back condition an average of 7.04 (SD = 2.24, range 1 – 15) occurred. In total, we sampled 7006 examples of experience in this study. In the current analysis, we calculated the mean scores of each question across the three sessions for each participant. The MDES scores were first transformed into z-scores for mean centring and unit variance scaling. The scores described the average momentary experience in each dimension. We use this score in the multivariate pattern analysis later.

**Table 1.** Multiple Dimension Experience Sampling questions in 0-back / 1-back task.

| **Dimensions** | **Questions** | **0** | **1** |
| --- | --- | --- | --- |
| **Task** | My thoughts were focused on the task I was performing. | Not at all | Completely |
| **Future** | My thoughts involved future events. | Not at all | Completely |
| **Past** | My thoughts involved past events. | Not at all | Completely |
| **Self** | My thoughts involved myself. | Not at all | Completely |
| **Other** | My thoughts involved other people. | Not at all | Completely |
| **Emotion** | The content of my thoughts was: | Negative | Positive |
| **Images** | My thoughts were in the form of images. | Not at all | Completely |
| **Words** | My thoughts were in the form of words. | Not at all | Completely |
| **Vivid** | My thoughts were vivid as if I was there. | Not at all | Completely |
| **Detailed** | My thoughts were detailed and specific. | Not at all | Completely |
| **Habit** | This thought has recurrent themes similar to those I have had before. | Not at all | Completely |
| **Evolving** | My thoughts tended to evolve in a series of steps. | Not at all | Completely |
| **Deliberate** | My thoughts were: | Spontaneous | Deliberate |

### 5.5 MULTIVARIATE PATTERN ANALYSIS

#### 5.5.1 Sparse canonical correlation analysis (SCCA)

We performed a sparse canonical correlation analysis (SCCA)^27,28^ on the functional connectomes and the cognitive tasks, to yield latent components that reflect multivariate patterns across neural organisation and cognition (For a similar application for experience sampling, see^29^). SCCA maximised the linear correlation between the low-rank projections of two sets of multivariate data sets with a sparse model to regularise the decomposition solutions a process that helps maximise the interpretability of the results. The regularisation function of choice is L_1_ penalty, which produces sparse coefficients, meaning that the canonical vectors (i.e., translating from full variables to a data matrix’s low-rank components of variation) will contain a number of exactly zero elements. L_1_ regularisation conducted (a) feature selection (i.e., select only relevant components) and (b) model estimation (i.e., determine what combination of components best disentangles the neurocognitive relationship) in an identical process. This way we handle adverse behaviours of classical linear models in high-dimensional data. A reliable and robust open-source implementation of the SCCA method was retrieved as R package from CRAN (PMA, penalized multivariate analysis, version 1.0.928). The amount of L_1_ penalty for the functional connectomes and cognitive task performance were chosen by cross-validation. The procedure is described in the section below.

Before SCCA, we regressed age, sex, head motion indicated by mean frame-wise displacement^53^ out of both the connectivity and cognitive task data to ensure that these potential confounders did not drive results. The implementation of the confound removal method was retrieved from Python library Nilearn (http://nilearn.github.io/, version 0.3.1). As features that do not vary across subjects cannot be predictive of individual differences, we limited our analysis of connectivity data to the top 5% most variable connections, as measured by median absolute deviation, which is more robust against outliers than standard deviation^54^. Thus the input data to the subsequent analysis consist of 247 ROI-ROI connectivity and 13 cognitive task measures.

#### 5.5.2 Model Selection

The model selection process was conducted in two parts: L_1_ penalty coefficient selection and component selection. For the L_1_ penalty coefficient selection, we performed a grid search combined with cross-validation (CV) to avoid over-fitting^55^. Of each penalty pair on the search grid, 5-fold cross-validation was performed to search for the best out-of-sample the rank-1 canonical correlation. We then decomposed the full dataset with the selected L_1_ penalty coefficients (see the top panel of Figure 6). The K-Fold CV was conducted by the implementation in Python library scikit-learn (http://scikit-learn.org/stable/, version 0.18.2).

**Figure 5.**
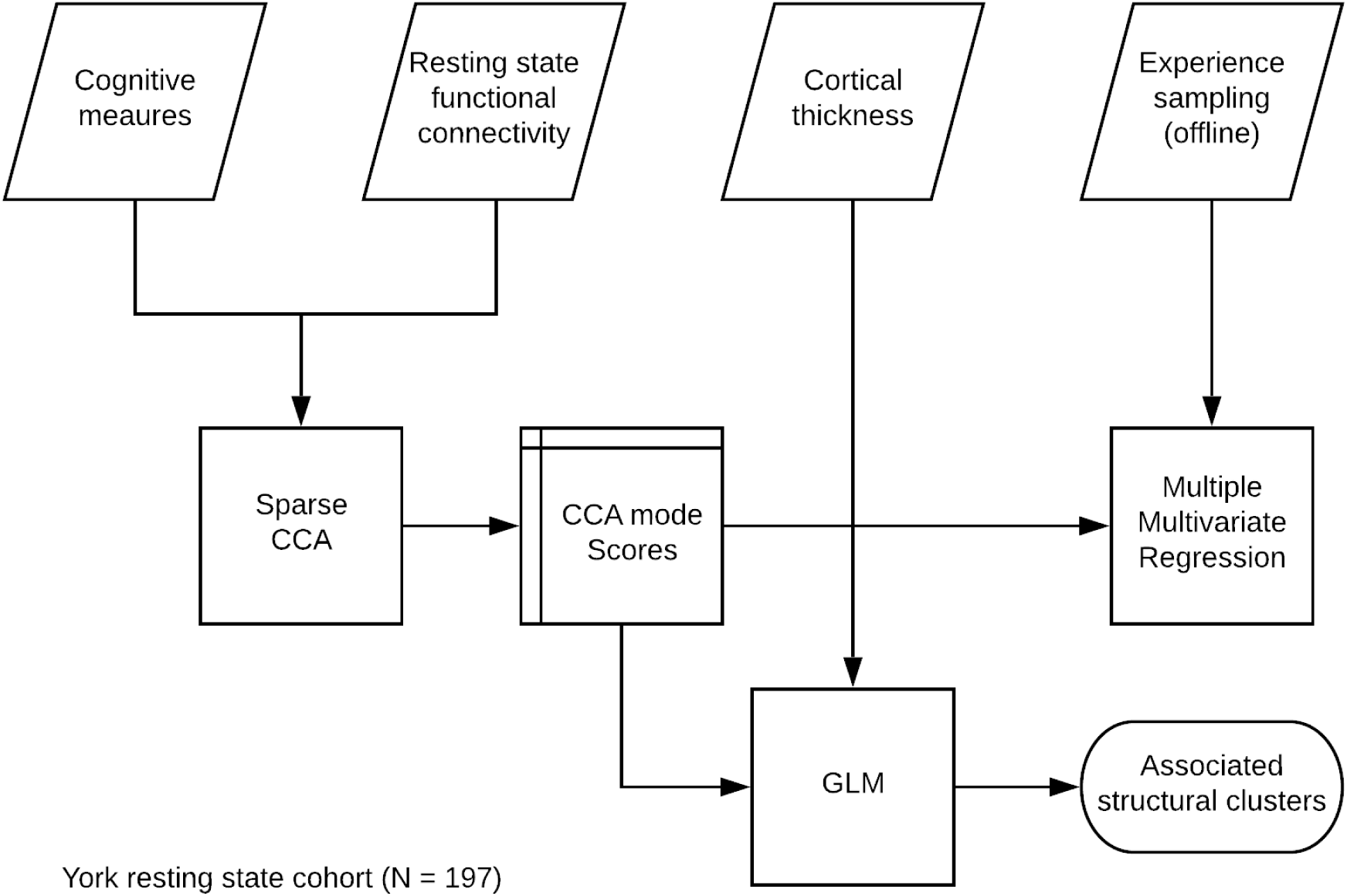
Analysis flowchart. The flowchart of the analysis pipeline. We first conducted SCCA to uncover the hidden structure that combines the task measures and the functional connectivity data. The SCCA model selection is detailed in 5.5.2 Model Selection and Figure 6. The theoretical validity of latent variables was later examined by predicting the ongoing thought report. We also explore the associated cortical thickness change associated with the functional neurocognitive modes. For the cortical thickness analysis, please refer to 5.6 cortical thickness analysis. For the details of multiple multivariate regression, please see 5.5.3 Group-level analysis.

**Figure 6.**
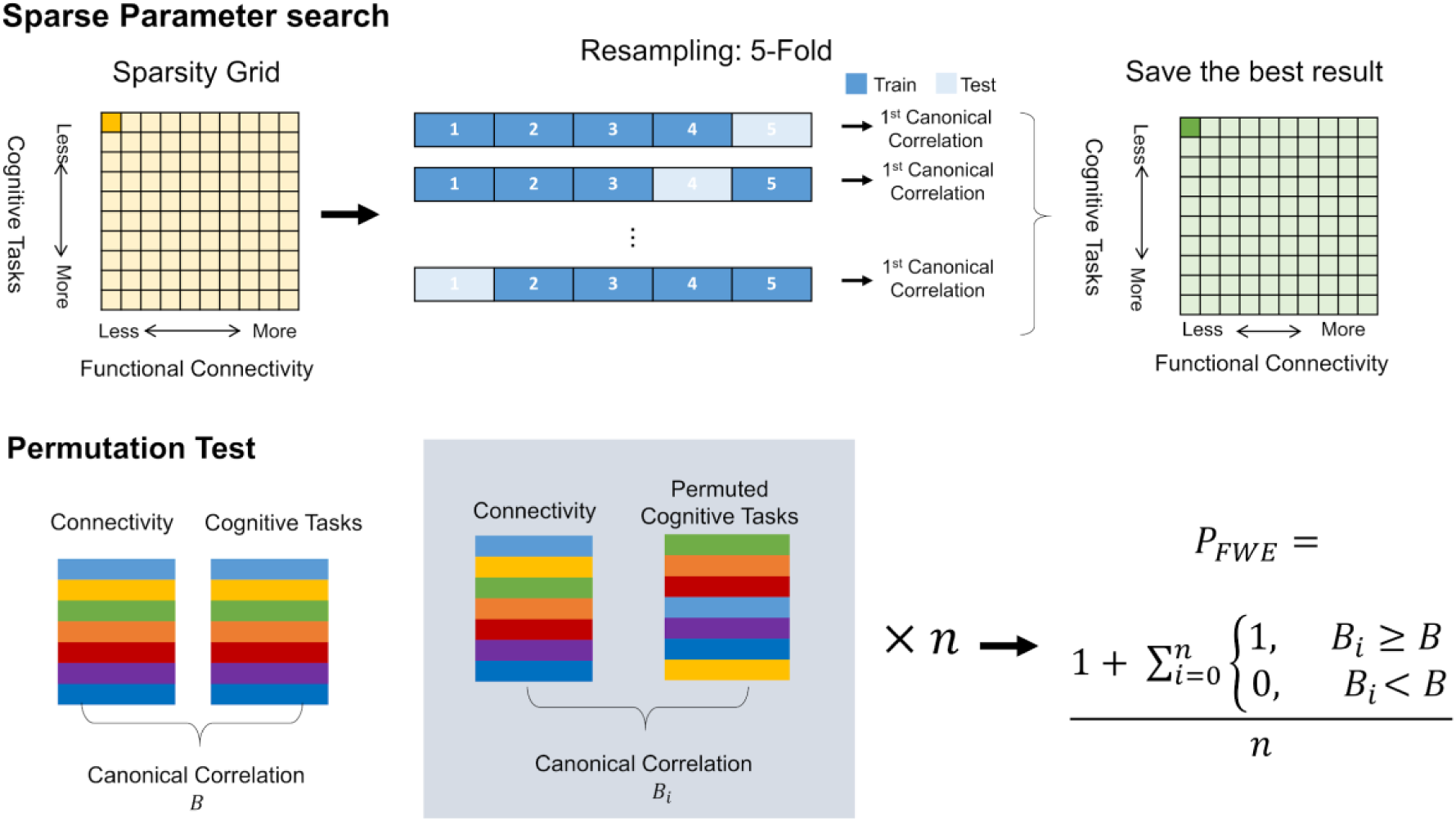
Multivariate pattern analysis pipeline. Top: parameter tuning combined with k-fold cross-validation to search for the best sparsity constraint with maximal out-of-sample the rank-1 canonical correlation. The selected set of parameters was then used as a basis to recompute CCA on the full dataset. Bottom: Permutation test for mode selection.

We performed SCCA with the optimal hyper-parameters on the full data set and saved all decompositions. A thousand permutations were applied to determine the mode(s) that occur above chance (see the bottom panel of Figure 6). We constructed an empirical null distribution for the set of canonical components by holding the functional connectivity data in place and permuting the row order of self-reports data. The permutation scheme specifically disturbed the link between individual differences in the dataset. The emerging distribution of no-association effects in our analysis setup provided the basis for testing the robustness of the components in the hypothetical population. A p-value with family-wise error (FWE) correction was calculated based on the permutation results. Modes are accepted with a 5% level of significance. In the current analyses, we adopt the permutation test with the FWE-corrected p-value by Smith and colleagues^56^. All modes were compared to the first canonical correlation of the permuted sample. The low-rank components are more relevant than the rest, therefore we yield more conservative p-value by comparing to the first canonical correlation only.

#### 5.5.3 Group level regression analysis

To determine how patterns of unconstrained neurocognitive activity related to performance on the ongoing thoughts, self-report experience summarised in three different ways (Overall average, average by days, and average by task), we conducted an independent statistical analysis on the identical subjects. The neuro-cognitive scores were calculated from averaging the canonical variates in the significant modes. A Type III multiple multivariate regression with Pillai’s trace test was applied to the data. Each of the latent components describing the neurocognitive mechanism from the SCCA was the independent variables, and the 13 measures from MDES were the dependent variables. We thus asked how well variability in the MDES items can be explained our robust components of brain-behaviour association across individuals. We hoped to describe the neuro-cognitive components by the linear combination of the self-report questions collected via MDES. The analysis was conducted in Python library Statsmodels (https://www.statsmodels.org/stable/index.html).

### 5.6 CORTICAL THICKNESS ANALYSIS

#### 5.6.1 Cortical thickness calculation

Freesurfer was used to estimate vertex-wise cortical thickness (5.3.0; https://surfer.nmr.mgh.harvard.edu). Briefly, the following processing steps were applied: intensity normalisation, removal of non-brain tissue, tissue classification and surface extraction. Cortical surfaces were visually inspected and corrected if necessary. Cortical thickness was calculated as the closest distance between the grey/white matter boundary and pial surface at each vertex across the entire cortex. A surface-based smoothing with a full-width at half maximum (FWHM) = 20 mm was applied. Surface alignment based on curvature to an average spherical representation, fsaverage5, was used to improve correspondence of measurement locations among subjects.

#### 5.6.2 Cortical thickness analysis

We explore the relationship between the SCCA modes (i.e. Their canonical variates) and cortical thickness to understand the structural effect of the neurocognitive components. The surfstat toolbox for Matlab^57^ was used for structural covariance network analysis, as in previous studies^58,59^. There is a well-established negative correlation between age and cortical thickness^60^ and gender also influences cortical thickness^61^. Consequently, these variables were included as covariates of no interest. The model fitted at a surface point i is shown as bellow: 

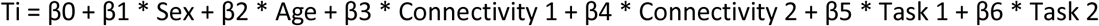

We determined significant clusters in this model using random field theory for nonisotropic images^62^ which controlled the Family-Wise Error rate at p < 0.05. We also explored the effect of the neurocognitive score. 

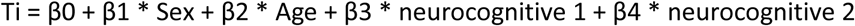

### 5.7 NEUROSYNTH META-ANALYSIS

We used the neurosynth meta-analytic database (https://www.neurosynth.org) to assess topic terms associated with the functional connectivity profile and significant cortical thickness regions. For each region of interest map, the output of the analysis was a z statistic associated with the feature term. The terms were then visualised as word clouds. Feature terms were derived from the 50 topic terms (v4). Of the 50, 25 were removed as “noise” terms or experiment paradigm related information, because they did not capture any coherent cognitive function, leaving 24 topic terms.

## 6 ACKNOWLEDGEMENT

This study was supported by the European Research Council (Project ID: 646927) (https://cordis.europa.eu/project/rcn/198057/factsheet/en) awarded to JS. The funders had no role in study design, data collection and analysis, decision to publish, or preparation of the manuscript. The authors extend their gratitude to Theodoros Karapanagiotidis, Deniz Vatansever, Charlotte Murphy, Mladen Sormaz, and Giulia Poerio for their invaluable contribution to the scanning of participants and data management. We thank Xiuyi Wag for the preprocessing of cortical thickness data. In addition, the authors thank the York Neuroimaging Centre staff for their support in setting up the imaging protocol and scanning. Finally, we thank all the participants for their time and effort in taking part in this study. The authors declare no conflict of interest.

## 6.2 AUTHOR CONTRIBUTIONS

HTW, EJ and JS designed the task and conceive the idea of the analysis. JS, DSM and EJ contributed to the development of the project. HTW analysed the functional and behavioural data. NSPH analysed the structural data. HTW, EJ and JS interpreted results. HTW and JS wrote the manuscript with comments from BCB, DB and EJ.

## Footnotes

* In the original study by Whitmer and colleagues^43^, contrast “switch cost”, smaller values indicates better ability to switch away from the previous condition. For the ease of interpretation, we reversed the scores and re-named the contrast as “flexibility”.

